# ClonoCluster: a method for using clonal origin to inform transcriptome clustering

**DOI:** 10.1101/2022.02.11.480077

**Authors:** LP Richman, Y Goyal, CL Jiang, A Raj

## Abstract

Clustering cells based on their high dimensional profiles is an important data reduction process by which researchers infer distinct categories of cellular state. The advent of cellular barcoding, however, provides an alternative means by which to group cells: by their clonal origin. We developed ClonoCluster, a computational method that combines both clone and transcriptome information to create hybrid clusters that weight both kinds of data with a tunable parameter. We generated hybrid clusters across six independent datasets and found that ClonoCluster generated qualitatively different clusters in all cases. The markers of these hybrid clusters were different but had equivalent fidelity to transcriptome-only clusters. The genes most strongly associated with the rearrangements in hybrid clusters were ribosomal function and extracellular matrix genes. We also developed the complementary tool Warp Factor that incorporates clone information in popular 2D visualization techniques like UMAP. Integrating ClonoCluster and Warp Factor revealed biologically relevant markers of cell identity.

## Introduction

Since the advent of high dimensional molecular profiling, clustering has been the most common form of data analysis and visualization applied, allowing one to form groups out of entities with similar profiles (Alon et al., 1999; Eisen et al., 1998). Clustering has enabled the detection of networks of disease genes and loss of function patterns across cancer cell lines (Cowley et al., 2014; Tripathi et al., 2019). More recently, the development of single cell measurement technologies has allowed the high dimensional profiling of individual cells. In this context, clustering has been used to categorize cells into discrete molecular states, often termed “fates” or “types”, that can be associated with distinct cellular functions (Kiselev et al., 2019; Peyvandipour et al., 2020). At the same time, in many biological contexts, single cells also have lineage relationships; i.e., they may arise from a common progenitor. This information in principle provides a complementary way to cluster profiles of single cells, but it has not been incorporated into the systematics of single cell profiles (Wagner and Klein, 2020).

Clustering purely based on molecular profiling has been successful in many instances but relies on a number of assumptions that have been difficult to evaluate rigorously. Briefly, most clustering approaches start with some form of feature selection or dimensionality reduction. Data points are then grouped to minimize the distances between elements of the same groups, either by k-means clustering, hierarchical clustering, or graph-based community detection methods that use connectivity to inform group membership (Kiselev et al., 2019; Menon, 2018; Peyvandipour et al., 2020). Intrinsic to all these methods is the use of some sort of distance between cells measured by some metric of distance between their molecular profiles, but in principle many other types of information could be used to inform or modify these distances.

Recently, the development of cellular barcoding systems has provided an alternative means by which to group cells. Primarily, these systems have been used for the longitudinal tracking of molecular profiles through some sort of biological process, such as differentiation or therapy resistance (Biddy et al., 2018; Emert et al., 2021; Fennell et al., 2022; Goyal et al., 2021; Oren et al., 2021; Umkehrer et al., 2021; Weinreb et al., 2020). Strikingly, these studies have concluded, in some instances, that single cell differences in the expression of few or even just one gene can predict the ultimate fate of the cell, differences that would not have been detected by standard clustering algorithms. For example, this approach identified the expression of *Mettl7a1,* a methyltransferase, as a driver of successful stem cell reprogramming (Biddy et al., 2018), identified TCF15 as necessary and sufficient for hematopoietic stem cell self-renewal, and identified *Pou2f2* expression in progenitor cells as predicting DC-like vs neutrophil-like monocyte fates (Rodriguez-Fraticelli et al., 2020; Weinreb and Klein, 2020; Weinreb et al., 2020). Such results suggest that the incorporation of clone information could be very useful in determining the expression differences between cells that have distinct functional outcomes. Being able to weigh both transcriptome and clone information in clustering algorithms would potentially enable the identification of such “hidden” factors.

We thus developed an algorithm that we call “ClonoCluster” that integrates transcriptome and clone barcode information, allowing one to cluster cells using a continuous parameter (α) that adjusts the relative weight of transcriptome vs. clone information. We applied ClonoCluster to six previously published independent single cell RNA sequencing datasets, including *in vitro* hematopoiesis, directed stem cell differentiation, and drug treatment of tumor cell lines (Goyal et al., 2021; Jiang et al., 2021; Weinreb et al., 2020). We found massive rearrangements of the assignments of cells to clusters as α was shifted to weigh clonal origin more heavily. These rearrangements had novel, potentially more biologically interpretable, cluster markers, and were associated with expression of genes involved in extracellular matrix production and translation. These results held across datasets where clone fate was determined intrinsically, and the effects were considerably less strong in the dataset where cell fate was determined extrinsically. Inspired by this clone-weighted network graph clustering approach, we developed a tunable parameter (the Warp Factor, ranging from 0-10), that incorporates clonality information into the dimensionality reduction step prior to the commonly-used UMAP algorithm for visualizing high dimensional datasets. We included ClonoCluster and Warp Factor into an open-source R package, *ClonoCluster* (https://github.com/leeprichman/ClonoCluster). As barcoding data becomes more prevalent, *ClonoCluster* can provide a means to evaluate the degree to which clustering can be altered by factoring in clonal origin.

## Results

### ClonoCluster integrates clone barcode and transcriptome information

Clone barcode assignment and transcriptome-level data represent two different modalities of data that can be used to cluster single cell RNA sequencing. In a prototypic clonal barcoding experiment, a population of cells is transfected with random transcribed barcodes such that each initial clone is likely to express unique barcodes. After proliferation, experimenters apply some additional experimental conditions, such as drug treatment or differentiation (Biddy et al., 2018; Emert et al., 2021; Goyal et al., 2021; Jiang et al., 2021; Weinreb et al., 2020). At the chosen endpoint, one can perform single cell RNA sequencing on a pool of individual cells, which is itself composed of some number of clones marked by barcodes. The barcodes themselves can be determined by various side reactions and subsequent sequencing, thus adding a clone identifier to each cell’s transcriptome. (In practice, technical constraints on clone identification and sampling for single cell RNA sequencing mean that only some subset of sequenced cells will have an identifiable barcode.)

Once cells have both transcriptome and clone information attached to them, one can compare methods of classification. Two popular software packages for classifying cells by transcriptome information alone are *Seurat* and *scanpy* (Satija et al., 2015; Wolf et al., 2018), both of which apply community detection algorithms to network graphs to identify the most interconnected cell clusters. We can then directly compare the classification of cells by transcriptome clusters vs. clone barcodes (Figure 1A). In principle, these two classification schemes could be virtually identical, or they could be completely uncorrelated with each other.

**Figure 1:**
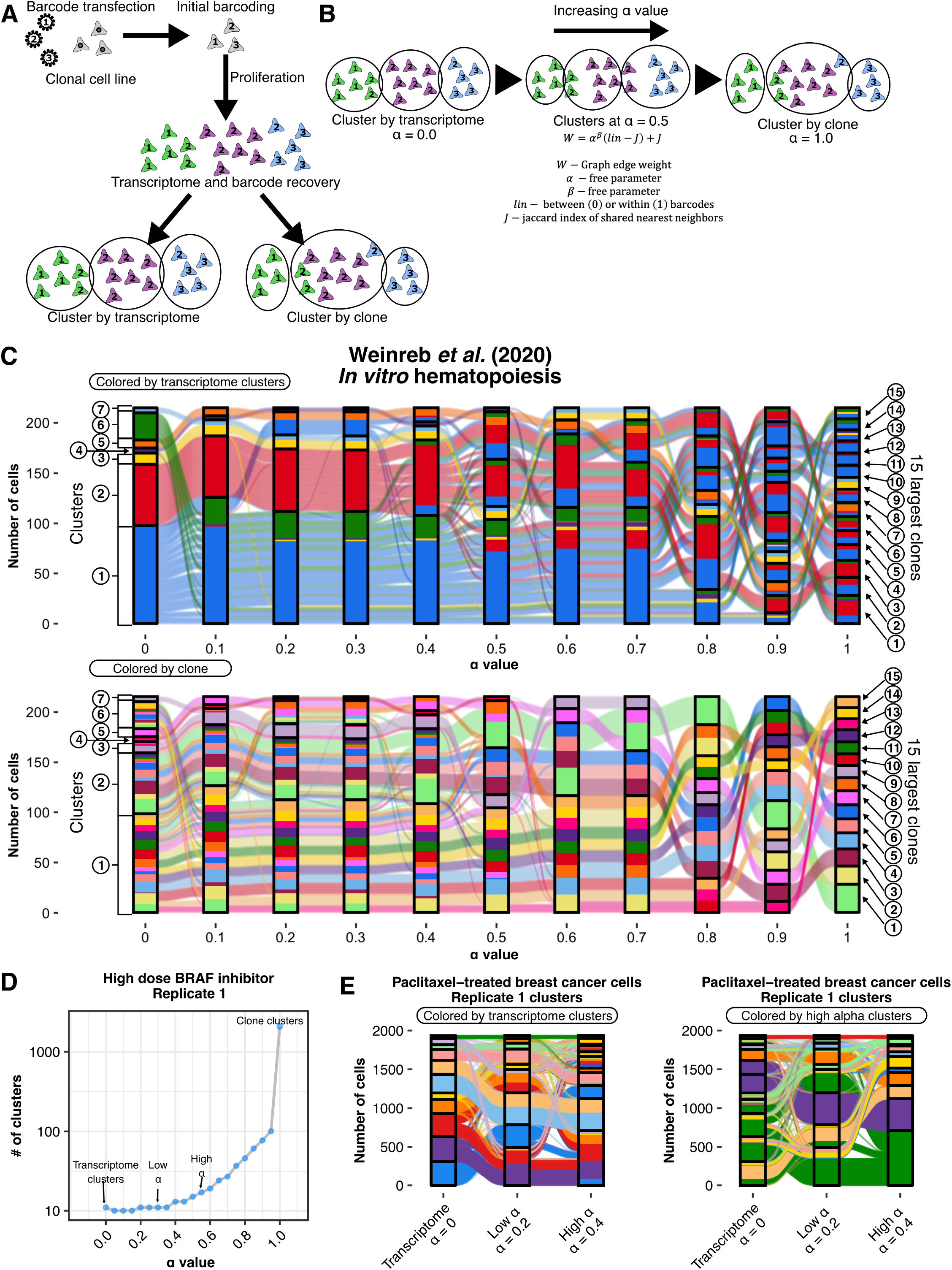
The ClonoCluster method integrates transcriptome and clone clustering modalities in a tunable manner using the α parameter. **A.** Schematic depicting a generic approach to single cell barcoding which yields output clusterable by two modalities of data, transcriptome clustering and clone clustering by recovered barcode. **B.** Schematic depicting the integration of these clone and transcriptome clustering modalities using the ClonoCluster method, in which the transcriptome nearest neighbor network graph edge weights are modified to incorporate clone clusters with a tunable free parameter, α. At α = 0, clustering is identical to traditional transcriptome clusters. At α = 1, clusters are consistent with clone barcode assignments. **C.** Sankey diagram depicting the reorganization of cell clusters from the 15 largest clone clusters present at day 2 in an *in vitro* hematopoiesis assay (Weinreb et al., 2020) with increasing α value colored by initial transcriptome cluster (top) and clone (bottom). Nodes/boxes represent clusters and ribbons depict the flow of cells between clusters. **D.** Representative plot of high dose BRAF inhibitor treated melanoma clonal cell line (clonal WM989 cells treated with 1μM of the BRAF inhibitor vemurafenib (Goyal et al., 2021)) showing that cluster number approaches unique clone barcode number with increasing α value at fixed community detection resolution. “High α” values were chosen based on the approximate shoulder of the curve after which clusters rapidly break into individual clone clusters with increasing α. “Low α” values were determined by the half “high α” value rounded to the nearest tenth value. At α = 0, clustering is identical to traditional transcriptome clusters. At α = 1, clusters are consistent with clone barcode assignments. **E.** Representative sankey diagram of the 15 largest clone clusters from the high dose BRAF inhibitor dataset treated melanoma clonal cell line depicting rearrangements of clusters at transcriptome, low α, and high α levels, colored by initial transcriptome cluster assignment (left) and high α cluster assignment (right).

### The tunable parameter α yields hybrid clone-transcriptome defined clusters

We wondered whether there was some way to incorporate both clone and transcriptome information to generate “hybrid” clusters that group cells that balance transcriptomic similarity and clonal relationships. In order to generate such hybrid clusters, we developed the ClonoCluster model for measuring similarity between cells. This model includes a tunable parameter, α, that shifts weight between clustering by cell transcriptomes alone (α = 0) and “clustering” by clone barcode alone (α = 1). In well-established single cell RNA sequencing analysis packages like *scanpy* and *Seurat,* the algorithms build a network graph of cells (nodes) connected by edges that are weighted by transcriptional similarity (“transcriptome weight”), determined by the number of shared nearest neighbors in principal component space (Satija et al., 2015; Wolf et al., 2018). The clustering itself is then determined by community detection within this graph, returning the most highly interconnected groupings of cells as the assigned clusters. In ClonoCluster, we retained this overall structure, incorporating clone information by modifying the weights as follows. For each edge between cells, we also created a “clone weight” of 1 or 0 based on whether the cells have the same or different barcodes. We then normalized the “clone weight” by the number of cells with that barcode to ensure that the attractive “force” of barcodes did not scale with the number of cells. We linearly combined transcriptome and clone weights using α such that it returned purely transcriptome weights for α = 0 and purely clone weights for α = 1 (Figure S1A). We could then use this graph for cluster assignment just as is done by the conventional algorithms. Thus, for values of α in between 1 and 0, ClonoCluster provides hybrid clusters that weigh both clone and transcriptome information.

### Clone to cluster concordance variably increases with α

We used ClonoCluster to generate hybrid clusters from six different clone barcoded single cell RNA sequencing datasets from our lab and others (Goyal et al., 2021; Jiang et al., 2021; Weinreb et al., 2020) (see Methods and Table 1). We first performed hybrid clustering at stepwise values of α from traditional transcriptome-only clusters (α = 0) to the clone groupings (α = 1). We wanted to determine how distinct hybrid clusters were from transcriptome or clone clustering alone as well as how increasing α reorganized clusters from initial transcriptome clusters to individual clone clusters. In order to visualize the flow of cells through these progressive hybrid clusters, we constructed Sankey diagrams (Kennedy and Sankey, 1898) of the datasets using the 15 largest clone clusters per sample (Figure 1C and S2A). There was visible reorganization of cells throughout clusters amongst the top 15 clone clusters as α increased, with differences in cluster number, composition, and size throughout much of the range of α in all samples. As expected, when α approached 1, the concordance between hybrid clusters and clone clusters increased, meaning that hybrid clusters were largely composed of individual clones or combinations thereof, with the frequency of clones being split across clusters becoming less. (At α = 1, clusters were equivalent to clone groupings.) Within each dataset, cells with the same clonal origin (i.e., sharing a barcode) demonstrated variable degrees of reorganization into concordant clusters at given levels of α, with some requiring higher α values to completely unify into a single hybrid cluster than others.

**Table 1:** Dataset descriptions and sources

| Data set name | Source | Description | # of Replicates | Fate determination |
| --- | --- | --- | --- | --- |
| Low dose BRAF inhibitor | Goyal *et al. biorXiv* (2021) | WM989 A6-G3 clonal melanoma cells treated with 100nM vemurafenib for 3-4 weeks | 1 | Intrinsic |
| High dose BRAF inhibitor | Goyal *et al. biorXiv* (2021) | WM989 A6-G3 clonal melanoma cells treated with 1μM vemurafenib 3-4 weeks | 2 | Intrinsic |
| Cario-directed iPSCs | Jiang *et al. biorXiv* (2021) | PENN123i-SV20 human IPSC line directed toward cardiomyocyte fate sequenced on day 14 | 1 | Extrinsic |
| *In vitro* hematopoiesis | Weinreb *et al. Science* (2020) | Hematopoietic progenitor cells derived from murine bone marrow differentiating in culture for two days | 1 | Intrinsic |
| WM983B BRAF inhibitor | Goyal *et al. biorXiv* (2021) | WM983B E6-C6 clonal melanoma cells treated with 100nM vemurafenib for 3-4 weeks | 2 | Intrinsic |
| Paclitaxel-treated breast cancer cells | Goyal *et al. biorXiv* (2021) | MDA-MB-231-D4 clonal breast cancer cells treated with 1nM paclitaxel for 3-4 weeks | 2 | Intrinsic |

The patterns of rearrangement also varied between datasets. For example, the cardio-directed iPSC dataset showed less concordance between clone clusters and transcriptome clusters amongst the top 15 largest clone clusters compared to WM989 A6-G3 melanoma cells treated with high dose vemurafenib. In the cardio-directed iPSC dataset, each of the top clone clusters segregated to a separate cluster by α = 0.6, compared to α = 0.8 or 0.9 in the two melanoma replicates (Figure S2A). We formally evaluated clone-to-cluster concordance using Cohen’s κ, a measure of agreement between classifications (grouping by clone vs. grouping by assigned cluster) computed from the 2×2 confusion matrix of clone barcode and cluster membership. When clonal cells sharing a barcode are more frequently assigned to a single cluster, Cohen’s κ is high. (At α = 1 where hybrid clusters and clonal clusters are equivalent, Cohen’s κ must also equal the maximum value of 1.) For most datasets, we observed a gradual transition of Cohen’s κ from 0 to 1 as α increased, suggesting that there was a graded conglomeration of clone barcodes within hybrid clusters as clonality was increasingly factored into the clustering algorithm. The cardiomyocyte-directed iPSCs, however, displayed a very sudden switch between low κ and high κ (Figure S2B), suggesting that there are few meaningful intermediate hybrid clusters between purely transcriptome clustering and purely clone clustering. Sankey visualization matched this interpretation by showing that clones were quite intermixed between the transcriptome-only clusters, with little obvious areas of partial conglomeration. In the cardiomyocyte-directed iPSC case, biologically, our prior findings suggested that cell fate was extrinsically determined and did not correlate well with clonality; hence, hybrid clusters would increasingly be composed of cells of essentially random types, explaining these observations for this dataset.

As α approaches 1, the number of hybrid clusters identified approaches the large number of individual clone clusters in the dataset, which are generally more numerous than the standard transcriptome clusters returned using common methods. We wondered if we could identify the maximum α value that generates a number of hybrid clusters similar to the number of transcriptome clusters that also showed significant reorganization of cells to be distinct from transcriptome-only clustering. We therefore computed the number of clusters returned in each sample across α values to identify the maximum α value that still returned a number of clusters in this range; the final cluster composition and cluster number was determined by the Louvain algorithm as implemented in the *Seurat* package (Blondel et al., 2008; Satija et al., 2015). The Louvain algorithm iteratively severs network graph edges to achieve optimal internal interconnectedness of community members (modularity) and returns a greater number of communities at higher values for the input resolution parameter. We performed all clustering at fixed resolution, therefore the rising number of clusters with increasing α value can be attributed to improved modularity of the clusters as networks break apart into isolated networks of like-barcoded clones. “High α” values were chosen based on the maximum value before the inflection point where cluster numbers dramatically increase, with “low α” values being chosen at half this value (Figure 1D and Figure S3). The high α value varied between samples, from 0.4 to 0.7. Even within this restricted range that limited the number of clusters returned, we observed reorganization at low and high α for the top 15 clone clusters (Figure 1E).

### Hybrid α clusters reveal novel cluster markers without loss of marker fidelity

A common goal of single cell RNA-seq analysis is the identification of “marker” genes whose expression is high in cells of a particular cluster (i.e., sensitive for cluster membership) and is low in cells in other clusters (i.e., specific for cluster membership) (Kiselev et al., 2019; Peyvandipour et al., 2020; Trapnell, 2015). Given that increasing α altered cluster number and membership, we wondered whether the genes that served as the best markers changed with α, and whether those markers were able to maintain similar sensitivity and specificity as they did on purely transcriptomically-defined clusters. We used the receiver operating characteristic (ROC) to evaluate all expressed genes as markers to classify cells into clusters at all possible marker expression cutoffs. Area-under-the-curve (which we refer to as “marker fidelity”) is derived from the ROC and summarizes the sensitivity and specificity across all marker thresholds, with an AUC of 0.5 indicating a classifier that is no better than random, and 1 being perfectly predictive. We identified the cluster markers with the highest AUC for each hybrid cluster at each α and found that the top markers of the clusters were different for different α values (Figure S4 and Figure S5A).

With this observed change in top markers, we wondered whether overall fidelity of the top cluster markers at each α, as reflected by AUC, was preserved with hybrid clustering. We found that across almost all datasets, the median AUC of top cluster markers at each value of α were not significantly different from transcriptome clustering alone (Figure 2A and Figure S5B), suggesting that the rearrangements caused by incorporating clone information did not radically decrease the ability to find a single marker for a cluster, despite the fact that the markers themselves changed. We did see decreased marker fidelity in the cardiomyocyte-directed iPSCs at the high α and clonal cluster levels (0.69 and 0.79 vs. 0.94). (Low dose vemurafenib-treated melanoma samples also had a decrease in marker fidelity, although to a lesser extent, 0.80 and 0.84 vs. 0.90.) Again, given the extrinsic determination in cardiomyocyte-directed iPSCs, we would expect that markers for hybrid clusters would be hard to find; indeed, many putative markers had poorer performance at high α (Jiang et al., 2021) (Figure 2D and Figure S4B). Thus, in the datasets analyzed in which cell states are thought to be intrinsically determined (Table 1), hybrid clustering yielded new cluster markers revealed by clonality information, often of equal fidelity to transcriptome clustering.

**Figure 2:**
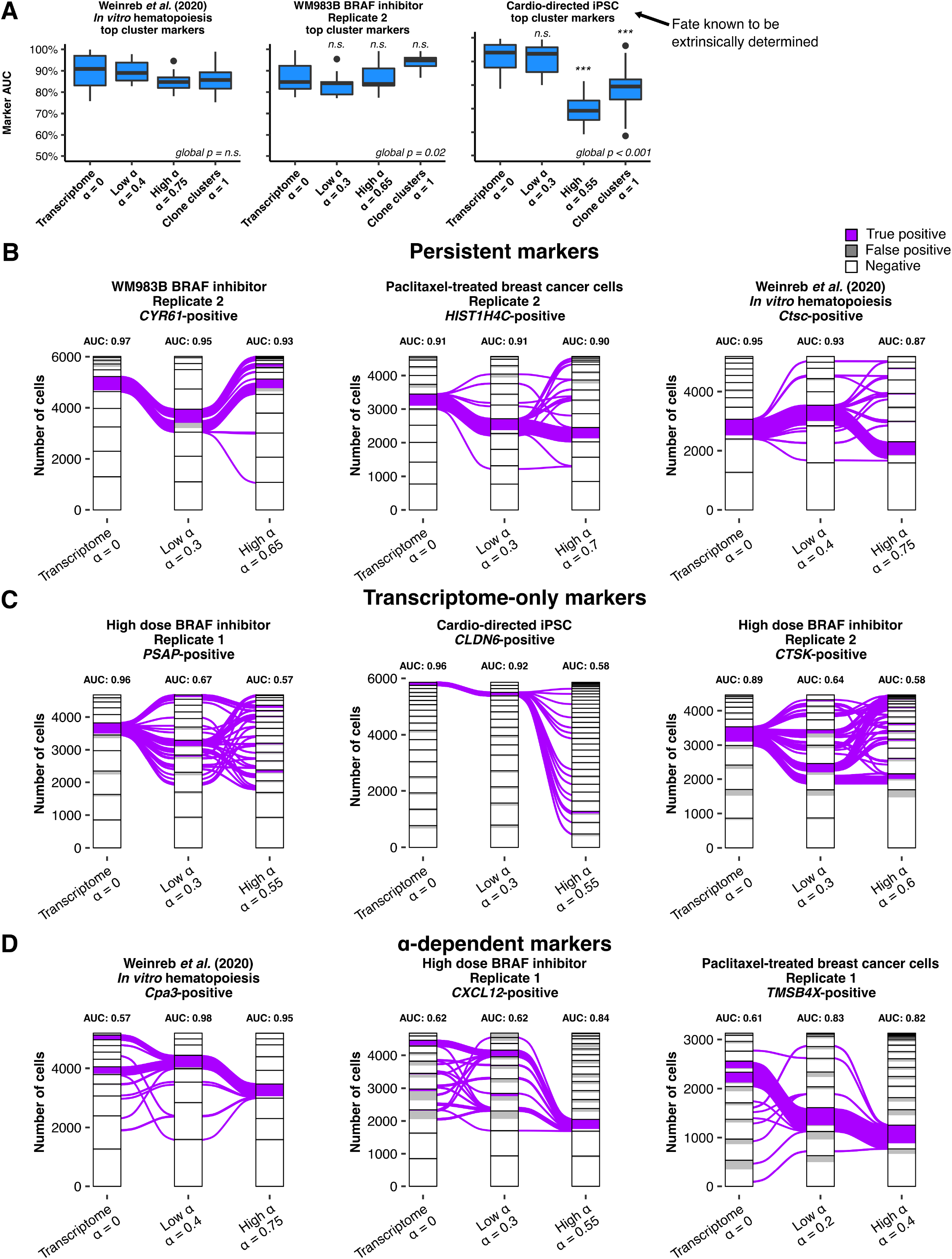
Manipulation of α reveals turnover of cluster markers **A.** AUC values for the top marker per cluster at zero, low, high, and maximum (α = 1) values of α for multiple samples, an *in vitro* murine hematopoiesis assay at day 2, a BRAF inhibitor treated clonal melanoma cell line WM989B, and directed differentiation of induced pluripotent stem cells (iPSCs) towards a cardiomyocyte fate, a system where extrinsic determinants of cell fate are expected be dominant over clone barcodes (Jiang et al., 2021). “Global p” indicates the p-value for the non-parametric Kruskal-Wallis test with Bonferroni correction. When “global p” was less than <0.05, pairwise Wilcox tests were performed with Bonferroni correction. Annotations above boxes indicate the corrected Wilcox test p-value compared to α = 0 (n.s. - not significant, *** - p < 0.001). **B – D.** Sankey diagram including all barcoded cells from various samples (see Methods) depicting transcriptome clusters, low α clusters, and high α clusters. Cells positive for the respective marker that are present in the cluster of interest are marked purple (“True Positive”) while cells positive for the marker but not present in the cluster of interest are marked gray (“False Positive”). Positivity thresholds were determined as described in Methods. Area-under-the-curve (AUC) for the marker is annotated above the cluster nodes. Representative markers are chosen to demonstrate markers whose classifier strength persists across α values **(A)**, markers that are only strong in transcriptome clusters **(B)**, and markers that are stronger at high and low α values **(C)**.

In addition to top cluster markers, we wondered how marker fidelity of any single marker was likely to change as we increased clonal weight with α. We used Sankey plots to directly visualize the flow of marker-positive cells through hybrid clusters as α changed. We found that the fidelity of transcriptome cluster markers varied as α changed, with some markers persisting across all values of α and others changing their fidelity dramatically. Many markers maintained their fidelity over increasing α values (representative samples shown in Figure 2A). We also identified markers that were strong cluster classifiers at the transcriptome cluster level that lost fidelity in low and high α clusters (Figure 2B), suggesting that the classification properties of these markers based on transcriptomes alone was lessened by the addition of clone information. Conversely, we also observed markers that increased in fidelity with increasing α, meaning that those markers were highly expressed in clones that were brought together into a hybrid cluster (Figure 2C).

### Reorganization of clusters with α is explained by differential expression of extracellular matrix and translation-associated genes

To identify expression of genes associated with reorganization of cells from transcriptome clusters to hybrid clusters, we evaluated genes as classifiers for explaining rearrangement of cells. Figure 3A demonstrates our approach to comparing reorganizing cells to delineate these differentially expressed genes. We have termed the area-under-the-curve for classifying a cell as one that will contribute to a hybrid α cluster of interest the reorganization AUC (“reorg-AUC”). Any gene with reorg-AUC greater than 0.80 for any transcriptome to hybrid α cluster was considered a “reorganization marker”. Representative examples of markers associated with contributing cells from a transcriptome cluster and a low α cluster of interest are shown in Figure 3B. Each set of reorganization markers for a paired hybrid cluster and contributing transcriptome clusters was then used to perform an overrepresentation analysis (Liao et al., 2019). Multiple gene sets were significantly enriched in reorganization markers from all samples. For visualization, we generated a heatmap of the maximum enrichment ratio for all gene sets significantly overrepresented in 3 or more samples, as well as several commonly chosen gene sets to serve as negative controls (Figure 3C). The gene sets shared across the greatest number of samples were associated with translation, “polysome”, “rRNA binding”, and “structural constituent of ribosome”, as well as a many samples showing enrichment for genes associated with the extracellular matrix, including “extracellular matrix”, “extracellular matrix structural constituent”, “extracellular matrix binding”, “collagen trimer”, “collagen binding”, and “fibronectin binding”. Enrichment was generally not seen in the negative control gene sets selected. From these analyses, it is evident that there are distinct biological underpinnings associated with cluster reorganization at even the low α level, including extracellular matrix and translation; remarkably, these results were consistent across independent datasets from very different biological processes.

**Figure 3:**
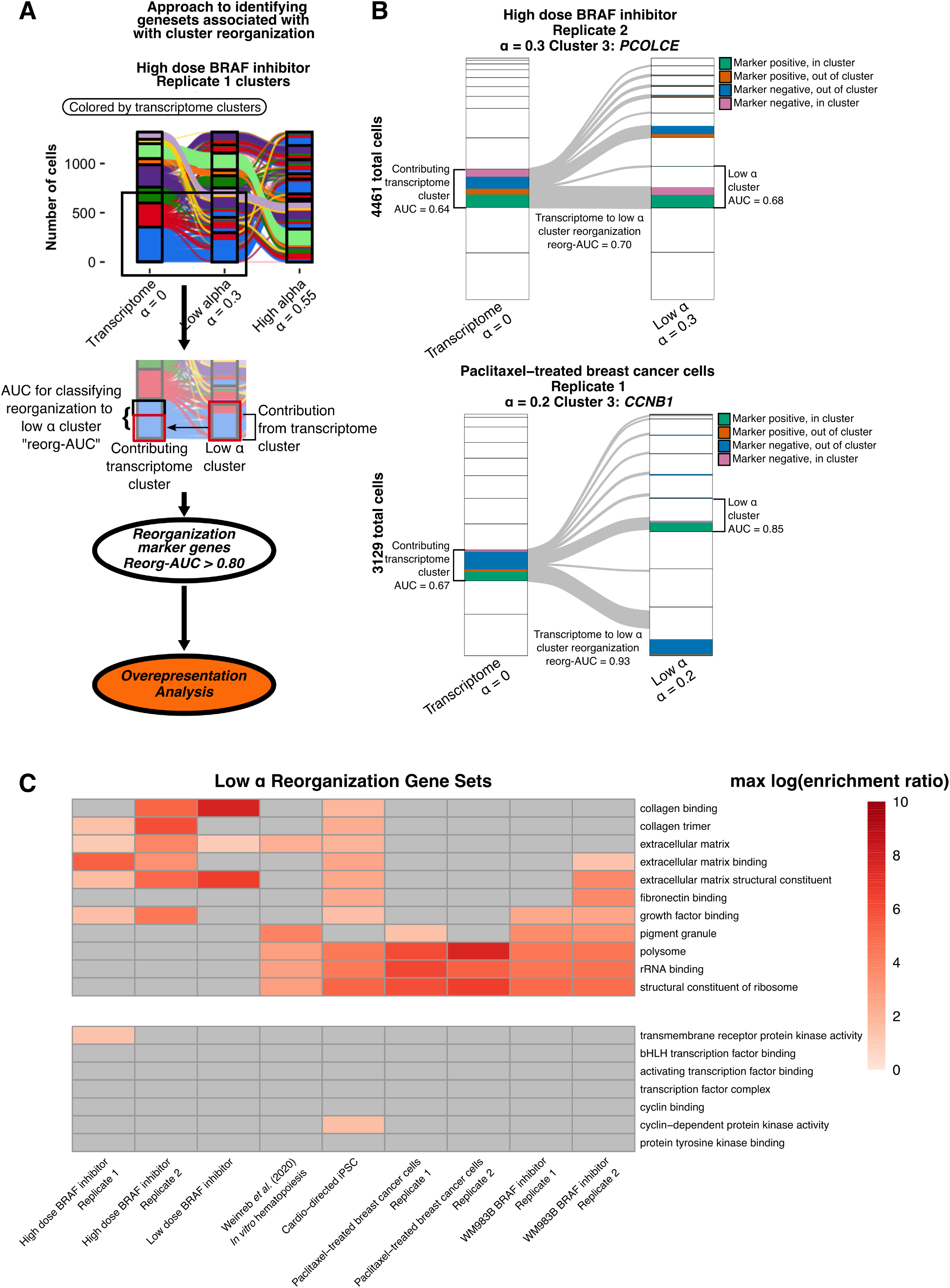
Reorganization markers are enriched in translation and extracellular matrix associated gene sets **A.** Schematic depicting approach to differential expression that explains cluster rearrangements with α modulation. For each hybrid α cluster and contributing transcriptome cluster pair, differential expression analysis is performed between cells inside and outside the α cluster, yielding an AUC for reorganization (reorg-AUC). Geneset overrepresentation analysis was then performed on these groups of differentially expressed genes on reorganizing cells with reorg-AUC > 0.80 (see Methods). **B.** Representative sankey diagrams showing a low α cluster and a contributing transcriptome cluster with colors indicating proportion of cells classified by the marker and the associated cluster AUCs and reorg-AUC. **C.** Heatmap demonstrating maximum log enrichment ratio for gene sets significantly enriched by overrepresentation analysis of reorganization markers across nine different data sources. All gene sets enriched in 3 or more samples are shown (top) as well as several commonly explored gene sets as negative controls (bottom). Gray tiles indicate no statistically significant enrichment in the sample (FDR > 0.05).

### Warp Factor, *s*, modifies UMAP representations to enhance clone separation

In addition to clustering and marker identification, an independent step in single cell RNA sequencing analysis is two-dimensional visual representation of transcriptome data using dimensionality reduction techniques such as t-SNE or UMAP (Becht et al., 2018; Kiselev et al., 2019). In the same spirit of hybrid clustering, we wondered whether we could apply the principle of adding weighted clone information to these low-dimensional visual embeddings. Whereas α modifies the weight of edges in the network graph input to the clustering algorithm, we sought to construct a method to incorporate clone information with a tunable parameter, the Warp Factor, into the input to the uniform manifold approximation and projection (UMAP) algorithm. UMAP represents variation in the transcript count matrix-derived principal component analysis (PCA) in a single manifold using attraction and repulsion components. Two UMAP dimensions are usually projected to visualize the high dimensional data (Becht et al., 2018; Kiselev et al., 2019). We modified the PCA input to UMAP to incorporate clone barcode information alongside transcriptome variation. To test this approach, we simulated data and used a model to create a modified principal component (PC) matrix in which a tunable “Warp Factor” parameter, *s*, warps the values of PCs for each cell to approach the mean value of the principal component for the clone cluster it belongs to (Figure S5A); i.e., reducing the variance of cells within the clone. At *s* = 0, the PC matrix is unmodified and at the maximum value, s = 10, the only variation present in the data will be between clone clusters (Figure S5B). The incorporation of the Warp Factor into visualization promoted separation of clone clusters in UMAP space in simulated data (Figure 4A). We then used the Warp Factor for the visualization of clone clusters in the datasets. As expected, the spread between individual clone clusters reduced across the UMAP with increasing Warp Factor. When we engaged high Warp Factors, individual clone clusters formed distinct spatial clusters in UMAP space. The amount of Warp Factor required to promote distinct separation of individual clone clusters and singlets into isolated spatial groups varied between datasets, likely due to the differences in the size and number of clone clusters as well as initial degree of clone-to-transcriptome concordance (Figure 4B). Unexpectedly, we also observed that “singlets”, cells that are the only member of their clone cluster (have a unique barcode), conglomerated together into a large group, possibly through repulsion from the non-singlet clone custers (Figure 4C). The Warp Factor method was effective at incorporating clonal information into UMAP visualizations of single cell RNA sequencing datasets by reducing within-clone cluster variation and increasing between-clone cluster variation in the underlying PC matrix.

**Figure 4:**
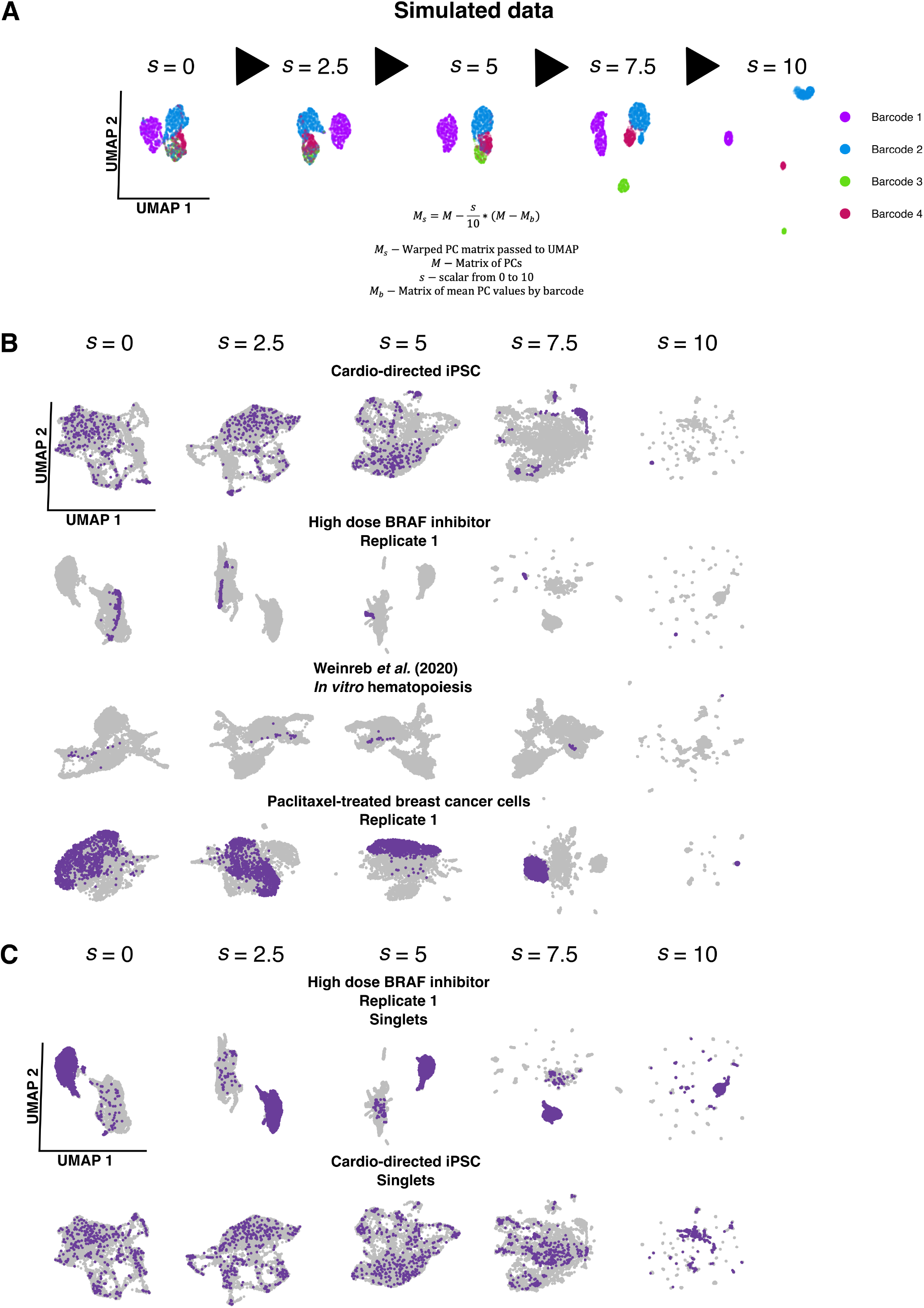
Warp factor, *s*, is a tunable parameter to modify UMAP visualization to incorporate clone barcode information. **A.** Demonstration of the effect of increasing Warp Factor (*s*) value on UMAP structure for simulated data of 3,000 cells with 4 barcodes and 6 principal components. Each UMAP axis is scaled and centered to allow comparison between facets. **B.** UMAPs with increasing Warp Factor for multiple datasets, highlighting a single large clone cluster in each **C.** UMAPs with increasing Warp Factor*s* showing singlets in two datasets. Singlets are cells that are the only in the sample with their unique barcode..

### Combining hybrid clustering and Warp Factor highlights unique clusters and markers

We sought to demonstrate the potential biological utility of ClonoCluster and Warp Factor when used together, focusing on data from melanoma (Goyal et al., 2021) and hematopoiesis (Weinreb et al., 2020). Starting with melanoma, we first identified top marker genes at a high α (α from 0.55 to 0.6); these markers were distinct from those identified by purely transcriptomic clustering (Figure 5A). Amongst these were *COL6A2* in low dose and *COL6A1* in high dose, which was consistent with our findings that collagen associated gene sets were enriched in reorganization markers (Figure 3C) in the three replicates of low and high dose vemurafenib-treated WM989 cells (Goyal et al., 2021). In the standard UMAP, cells that highly expressed these markers were dispersed, but upon engaging a Warp Factor of 5, these cells came together, demonstrating that Warp Factor can visually represent the results of incorporating clonal information.

**Figure 5:**
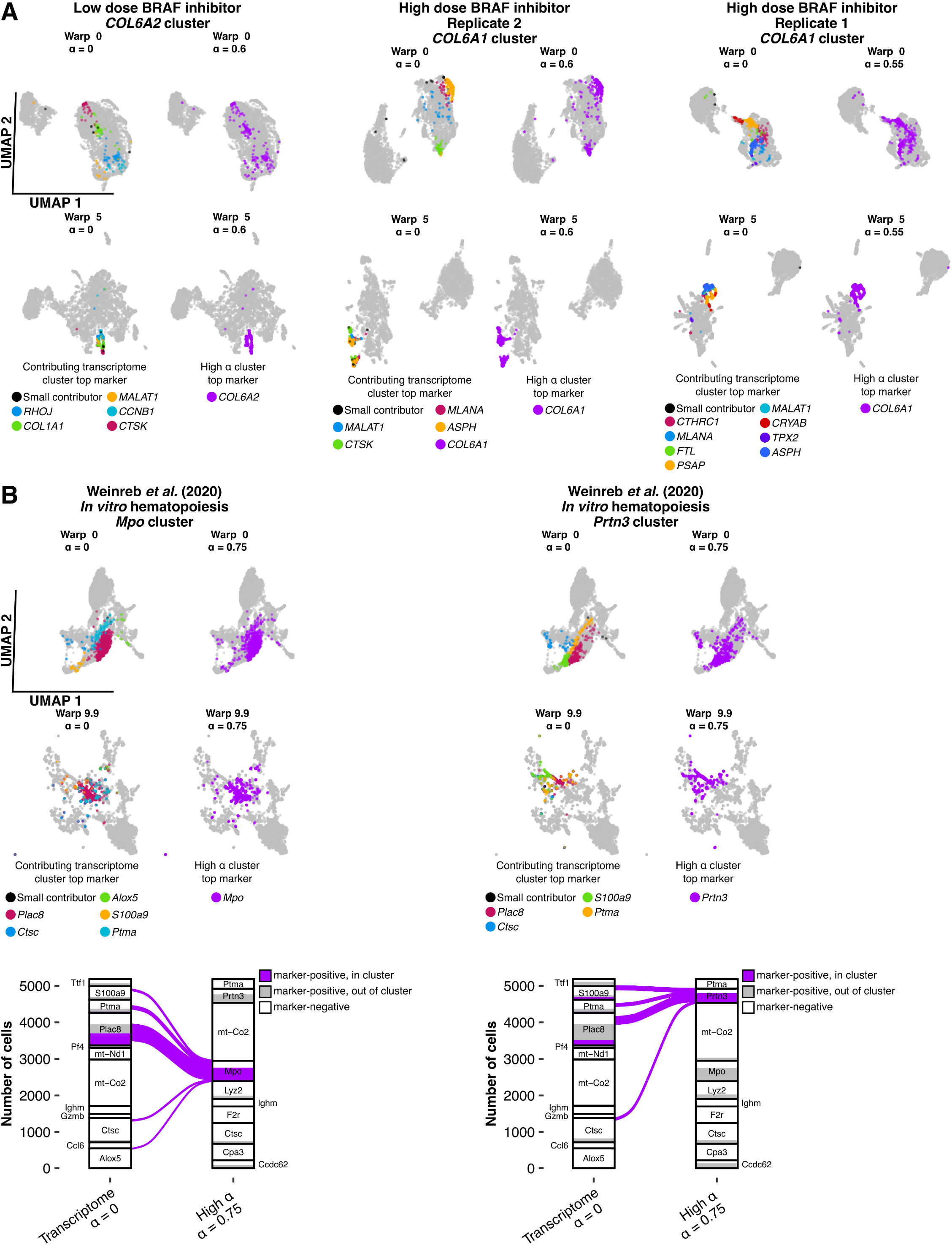
Combining hybrid clustering and Warp Factor highlights clusters with distinct markers in UMAP representations. A. UMAPs highlighting a single high α cluster with the indicated top marker gene *(COL6A1* and *COL6A2)* in different replicates and doses of WM989 cells treated with vemurafenib (Goyal et al., 2021). Representations are stratified by α and Warp Factor. Gray points represent cells outside the high α cluster. Colors reflect the top marker gene of the high α cluster (purple) or the contributing transcriptome clusters. Contributing transcriptome (α = 0) clusters were grouped as “small contributors” if less than 10 cells were present in the high α cluster of interest. B. UMAPS as in **(A)** for the day 2 *in vitro* hematopoiesis data (Weinreb et al., 2020) highlighting high α clusters marked by the granulocyte-enriched markers *Mpo* and *Prtn3* (top) and Sankey diagram depicting marker positivity at both clustering levels, nodes are annotated with top cluster markers. “Small contributors” indicates all contributing transcriptome clusters that contributed less than 10 cells to the high α cluster.

We also performed a similar evaluation on the day 2 *in vitro* hematopoiesis data from Weinreb *et al.* The original authors labeled the majority of these cells as “undifferentiated” at day 2, as they did not meet the expression level for conventional markers used for supervised cutoffs to define cell types (Weinreb et al., 2020). Indeed, using conventional transcriptome clustering, many of the top cluster markers identified *(Alox5, Plac8, S100a9, Ptma,* and *Ctsc)* were not ones known to be associated with distinct cell types in this system. However, for high α (α = 0.75), two markers emerged as top cluster markers *(Mpo* and *Prtn3)* that are known neutrophil-specific genes (Fig. 5B) (Kessenbrock et al., 2009). Using a Warp Factor of 9.9 (likely needed due to the small number of cells per clone), we could pull these markers together to some degree. The fact that ClonoCluster was able to recover more biologically meaningful markers suggests that there may be biological utility in its use.

## Discussion

Clonal barcoding has provided additional information that could be used for clustering cells based on both clonal origin and transcriptome. ClonoCluster provides a method for weighting clone and transcriptome information—using a tunable parameter α—to generate hybrid clusters. These clusters were distinct from purely transcriptomically-defined clusters, with unique sets of marker genes. The reorganization was often accompanied by the alignment of the expression of extracellular matrix proteins, suggesting that category of protein may be important for discriminating clones from each other. Further, we developed Warp Factor, inspired by ClonoCluster, as a way to modify the popular UMAP visualization to incorporate clone information.

A major question is whether these hybrid clusters more accurately reflect biological differences than transcriptomically defined clusters. We remain agnostic to this question, and pose ClonoCluster as a method to tune the degree to which clonal information is incorporated, with the degree of tuning left to the user to decide. Nevertheless, there are suggestions that some degree of clone information does reveal biological information. First, the fact that the assignment of cells to clusters changes dramatically as α is changed suggests that there is at least different information captured by clone information. Moreover, the top markers for clusters changed markedly, again suggesting different biological characterizations. It is hard to know what markers are biologically “correct”, but we do point out that in the case of the *in vitro* hematopoiesis dataset, we found that increased α let to the detection of *Mpo* and *Prtn3* as a markers, which are well-known neutrophil-specific genes (Kessenbrock et al., 2009) that did not appear as a top cluster marker for pure transcriptomic clustering. Moreover, in the case of cardiomyocyte-directed iPSCs from Jiang *et al.* (Jiang et al., 2021), in which we know that cell fate is largely determined by extrinsic factors and thus less by clonal factors, we observed that marker fidelity decreased at the high α levels, providing a negative biological control for ClonoCluster. Ultimately, further testing of the homogeneity in the functional properties of cells within each cluster will be required to truly determine what clustering methods most closely match biological distinctions.

What are the primary transcriptomic correlates with the rearrangements driven by incorporating clone information? We found these correlates were primarily associated with translation, ribosomal activity, and components of the extracellular matrix. These associations were found across diverse, independent datasets, suggesting that it is a common feature of transcriptomes of clones. There is a known association between ribosomal gene expression and overall gene expression level that may be the source of its influence on cell clustering (Kiselev et al., 2019). Extracellular matrix may indicate some memory of microenvironmental differences that differ between clones. Such memory would have to be encoded intrinsically, however, because “twin” experiments have shown that the same clone, exposed to two different microenvironments post-drug have virtually identical transcriptomes (Goyal et al., 2021). It is thus also possible that extracellular matrix proteins inherently encode important information about cell types (Sacher et al., 2021). It is interesting that there is a shift from more canonical markers of cell types, such as transcription factors, towards these other factors, suggesting that transcription factors may not be the sole determinants of cell type.

Technology for reading out full lineage data (as opposed to just clone data) has now been developed, often using CRISPR/Cas9 genome editing to make mutations in the genome that can be read out by sequencing (Frieda et al., 2017; Kalhor et al., 2017; Raj et al., 2018; Spanjaard et al., 2018) or imaging to generate the full cellular family tree (Chow et al., 2021; Frieda et al., 2017; Packer et al., 2019; Yu et al., 2009). It is relatively straightforward to adapt ClonoCluster to these situations, in which one can weight the distance between cells by the relative phylogenetic distance.

In general, the problem we describe here is in many ways analogous to whether species should be organized by genetic phylogeny or by phenotypic characteristics (Hull, 2010). While the debate within systematics has not yet been resolved, we believe that field may have much to inspire current research in categorization of cells based on molecular profiling.

## Methods

### Source data

Single cell RNA sequencing and barcode data for six datasets were compiled from multiple sources. “Low dose BRAF inhibitor” and “high dose BRAF inhibitor” indicate replicates of a monoclonal human melanoma cell line, WM989 A6-G3 (Emert et al., 2021), transfected with a barcode library and treated with 250nM and 1μM vemurafenib respectively. “WM983B BRAF inhibitor” samples indicates replicates of a vemurafenib-resistant monoclonal human melanoma cell line, WM983B E6-C6 (Emert et al., 2021) transfected with a barcode library and treated with 250nM vemurafenib (Goyal et al., 2021). “Paclitaxel-treated breast cancer cells” indicates replicates of a monoclonal line, MDA-MB-231-D4, derived from human breast cancer cell line transfected with a barcode library and treated with 1nM paclitaxel, as previously described (Goyal et al., 2021; Shaffer et al., 2020). “Cardio-directed iPSC” indicates a barcode-transfected induced pluripotent stem cell line treated to drive cells towards a cardiomyocyte fate as previously described (Jiang et al., 2021). Count matrices, barcoding methods, and barcode assignments for these samples were generated as described (Goyal et al., 2021; Jiang et al., 2021).

*“In vitro* hematopoiesis” samples indicate barcoded murine hematopoietic stem cells differentiating *in vitro,* collected and sequenced at the day 2 time point when most cells were labeled “undifferentiated” using known cell markers. Count matrices and barcode assignments were deposited by Weinreb *et al.* (2020) (Weinreb et al., 2020) and retrieved from the NCBI Gene Expression Omnibus at https://www.ncbi.nlm.nih.gov/geo/query/acc.cgi?acc=GSE140802. Count matrices and barcodes for all datasets used in this study are available in the analysis repository on GitHub (https://github.com/leeprichman/ClonoCluster_paper).

### Transcriptome and clone barcode integration with ClonoCluster

To integrate the transcriptome and clone barcode information, the shared nearest neighbors network graph is produced, using the single cell RNA sequencing analysis R package *Seurat* (Satija et al., 2015). Briefly, gene counts are converted into 100 principal components using the *irlba* package (Baglama et al., 2021). *k* nearest neighbors in PCA space are identified and a network graph is constructed where cells are nodes and edges between any two cells are given weight equivalent to the jaccard index between shared nearest neighbors (defined below as *J).* For all analyses, *k* was set to 20 and the graph was pruned of any edges with weight less than 1/15, consistent with *Seurat* defaults.

A size-normalized clone barcode network graph is also constructed where each cell is represented as a node, with edges drawn between all cells with weights of 0 for cells that are not in the same clone cluster (i.e. do not share a barcode) or 1/*n* for cells that are in the same clone cluster (i.e. do share a barcode), where *n* is equal to the total number of cells assigned to that barcode (size of the clone cluster.) We found that size normalization is necessary to prevent the highly interconnected larger clone clusters from exerting a dominant effect on graph modularity and final clustering compared to smaller clone clusters.

The transcriptome and clone graph edge weights are integrated using the following equation:

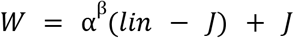

Where *W* is the final output edge weight between two cells, α is a chosen parameter ranging from 0 to 1, ß is a chosen parameter from 0 to 1, *lin* is the value of the edge in the clone barcode graph {0,1/*n*} and *J* represents the edge weight in the transcriptome graph: the jaccard index of shared nearest neighbors in transcriptome space. At α = 0, the edge weight between any two cells is therefore equivalent to the transcriptome graph (*J*), and at α = 1 the edge weight is equivalent entirely to the clone barcode graph (*lin*). ß modifies the effect of step sizes in α, lower values of ß increase the effect of α on final edge weight *W* at low α. For all datasets and analyses, ß was set to 0.1. The combined graph with modified edge weights is then passed to the Louvain community detection algorithm (Blondel et al., 2008) as implemented in *Seurat* to return cluster assignments.

### Cluster assignments

Resolution for the community detection algorithm was fixed for each sample, ranging between 0.6 and 1 across all nine replicates, with an initial target of 8 – 20 clusters at α = 0. As α approaches 1, the number of clusters returned at fixed resolution approaches *n*, the number of unique clone barcodes present. Therefore, the “high α” level was identified by approximation of the α value at the inflection point where the number of clusters rapidly increases and begins to reflect single clone clusters. The “low α” level was defined as half the “high α” value rounded to the nearest tenth. This gave the maximum number of clusters as 32 at the high α level in the Jiang *et al.* dataset.

### Clone barcode to cluster correlation analysis and visualization

Sankey diagrams were generated using the *ggalluvial* R package. For Sankey diagrams inclusive of α = 1, the top 15 largest clone clusters per sample are chosen for ease of visualization and limitation of discrete colors used. Unless otherwise specified, the complete dataset is shown for Sankey diagrams. Cohen’s κ was computed at each α level per clone barcode for the hybrid cluster with the largest proportion of the barcoded cells within it.

### Cluster marker identification

Cluster markers were identified by receiver operating characteristic (ROC) using the *ROCR* package (Sing et al., 2005). All genes present in the count matrix were tested for classification of a cell within a cluster versus all other clusters combined. When cells are noted as “marker positive”, the threshold for positivity was identified from the point on the ROC curve with minimum Euclidean distance from 100% TPR and 0% FPR.

### Reorganization marker and overrepresentation analysis

For each cluster of interest at a given α value, cells from each contributing cluster at the transcriptome level (α = 0) were compared to all other cells from the contributing transcriptome cluster. ROC analysis was performed to determine the strength of genes in the count matrix to classify cells from the contributing transcriptome cluster that reorganize to the α cluster of interest as long as the two comparison groups contained at least 10 cells. The AUC from this analysis was termed the reorganization AUC or reorg-AUC. Reorganization marker genes were defined as genes with reorg-AUC > 80. Each set of reorganization markers for the paired hybrid α cluster and contributing transcriptome (α = 0) cluster was used for overrepresentation analysis. When a transcriptome cluster contributed in its entirety to an α cluster (i.e. the transcriptome cluster was a subset of the α cluster), the AUC for overall cluster markers for the contributing transcriptome cluster was considered the reorg-AUC. Thresholds for marker positivity for visualization were determined from the point on the ROC curve with minimum Euclidean distance from 100% TPR and 0% FPR.

Overrepresentation analysis was performed with the R package *WebGestaltR* (Liao et al., 2019). The search space included the Gene Ontology Cellular Components and Molecular Function datasets (Ashburner et al., 2000). Significantly overrepresented gene sets were defined as those with FDR < 0.05. For visualization, heatmaps were created of the maximum log base 2 of the enrichment ratio for any cluster reorganization in the sample.

### Modified principal components and UMAP visualization

Uniform manifold approximation and projection (UMAP) is a dimensionality reduction technique frequently used to visualize gross trends in high dimensional single cell RNA sequencing data (Becht et al., 2018; McInnes et al., 2018). Clustering assignments and associated markers computed from principal component space are usually overlaid as well (Kiselev et al., 2019). To incorporate clone barcode information into the two dimensional UMAP representation of data, we modified the principal components using a tunable parameter *s* according to the following equation:

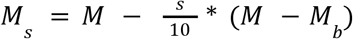

Where *M_s_* is the modified PCA matrix passed to the UMAP algorithm, *M* is the PCA matrix, *s*, Warp Factor, is a parameter from 0 to 10, and *M_b_* is a matrix composed of the average value of the PCA matrix per clone. At *s* = 0, the input to the UMAP will be the PCA matrix, *M*. At *s* = 10, the input to the UMAP will be a matrix with the value of each principal component substituted for the corresponding mean value per clone barcode, *M_b_*, thus the only variation reflected in the UMAP will be between clones (Figure S5B). The parameters for UMAP as implemented in the *uwot* package were identical to the default settings of *Seurat* (McInnes et al., 2018; Satija et al., 2015).

### Data visualization and statistical analysis

Data manipulation and statistical analysis was performed in the R statistical computing language with the *data.table, stats, ROCR, WebGestaltR,* and *magrittr* packages. Transcriptome nearest neighbor graph construction and community detection was performed with the *Seurat* package. Data visualization used the R packages *ggplot2, pheatmap, ggalluvial,* and *VennDiagram.* Global hypothesis testing was performed using the nonparametric Kruskal-Wallis test with Bonferroni correction for multiple comparisons. Local testing was performed with pairwise Wilcox tests with Bonferroni correction.

### Software availability

Raw data and scripts for all analyses are available at https://github.com/arjunrajlaboratory/ClonoCluster_paper. ClonoCluster is open source and available under the GPL3 license at https://github.com/leeprichman/ClonoCluster, including worked examples, example data, unit tests, and a Docker image.

---

**Figure S1:**
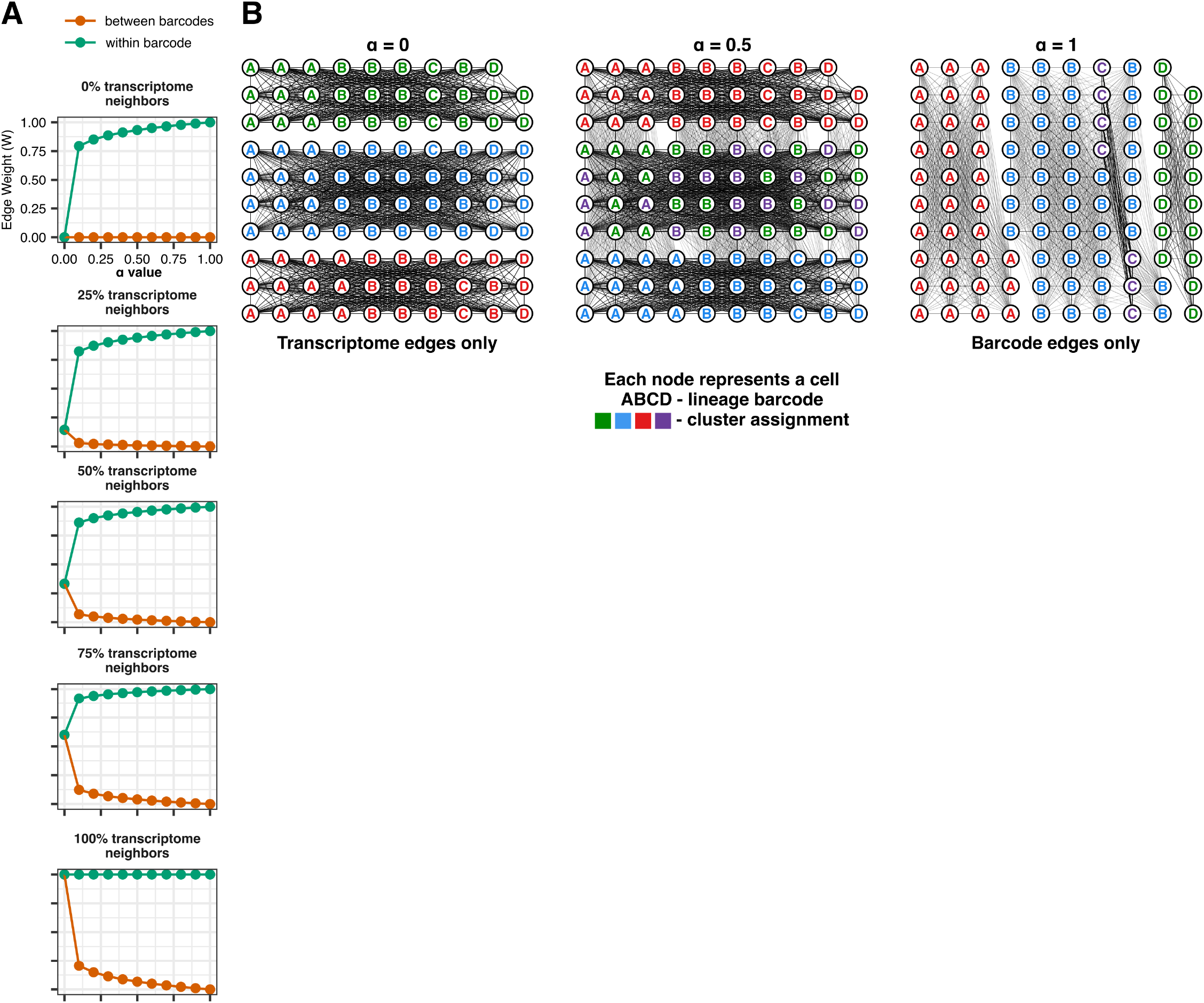
α modifies network graph edge weights to create Louvain clusters that increasingly reflect clone barcode assignments. A. Output edge weight of the ClonoCluster model for graph edge weights between two cells for different proportions of shared nearest neighbors in transcriptome space with fixed ß = 0.1. B. Network graphs with modified edge weights and cluster assignments at three α values for simulated data.

**Figure S2:**
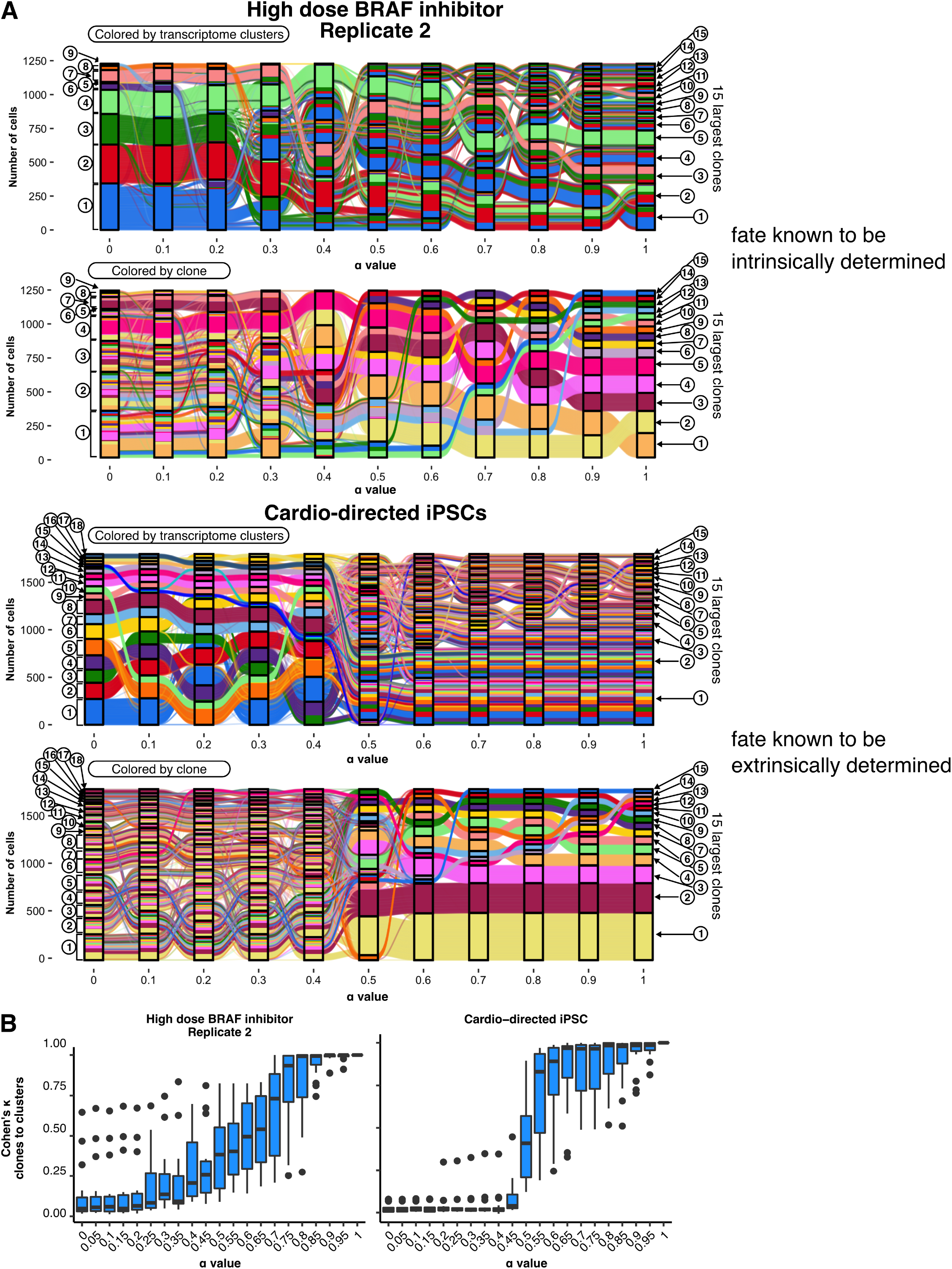
Intrinsic vs extrinsic determinants of cell fate stratify the effect of α on clone-cluster congruence. **A.** Sankey diagrams for the 15 largest clone clusters in two samples across α values, a clonal melanoma cell line treated with high dose (1μM) vemurafenib (Goyal et al., 2021) with known intrinsic determinants of cell fate and directed differentiation of induced pluripotent stem cells (iPSCs) towards a cardiomyocyte fate (Jiang et al., 2021) with previously described dominance of extrinsic determinants of cell fate. **B.** Cohen’s κ for interrater reliability for classification of a cell by clone barcode and cluster assignment for the top 15 largest clone clusters across α values for the samples in **(A)**.

**Figure S3:**
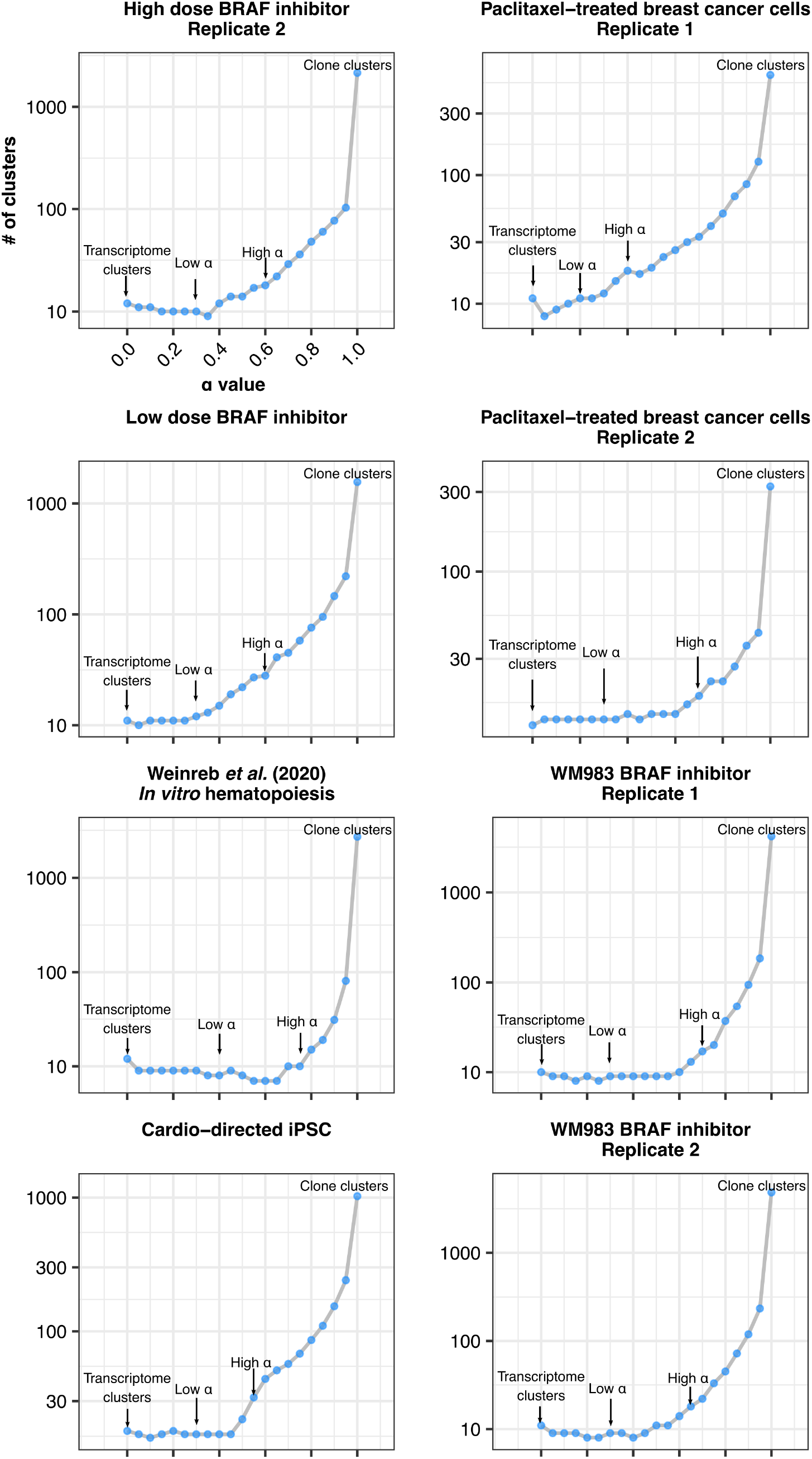
The number of clusters returned by community detection at fixed resolution approaches the number of unique clone barcodes in the data as α approaches 1. Plots showing the number of clusters returned by community detection at constant resolution with increasing α in each dataset. At α = 1, the network graph is composed of like-barcoded cells (a clone cluster) connected only to each other and community detection returns the number of unique clone clusters in the data as the number of clusters.

**Figure S4:**
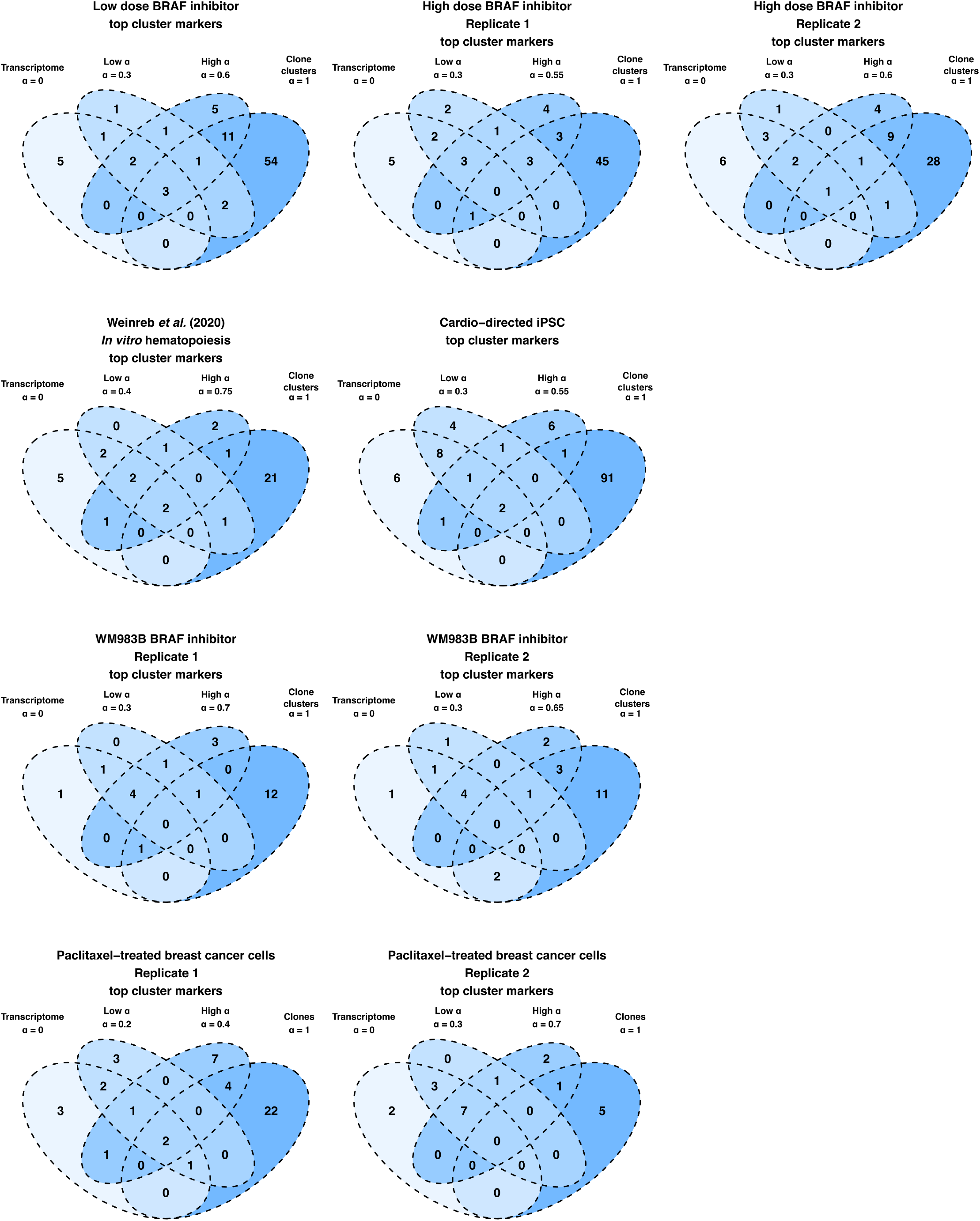
Significant marker turnover occurs between transcriptome, low α, high α, and clonal clusters. Venn diagrams showing the number and overlap of markers with AUC > 0.70 for any transcriptome, low α, high α or clone cluster.

**Figure S5:**
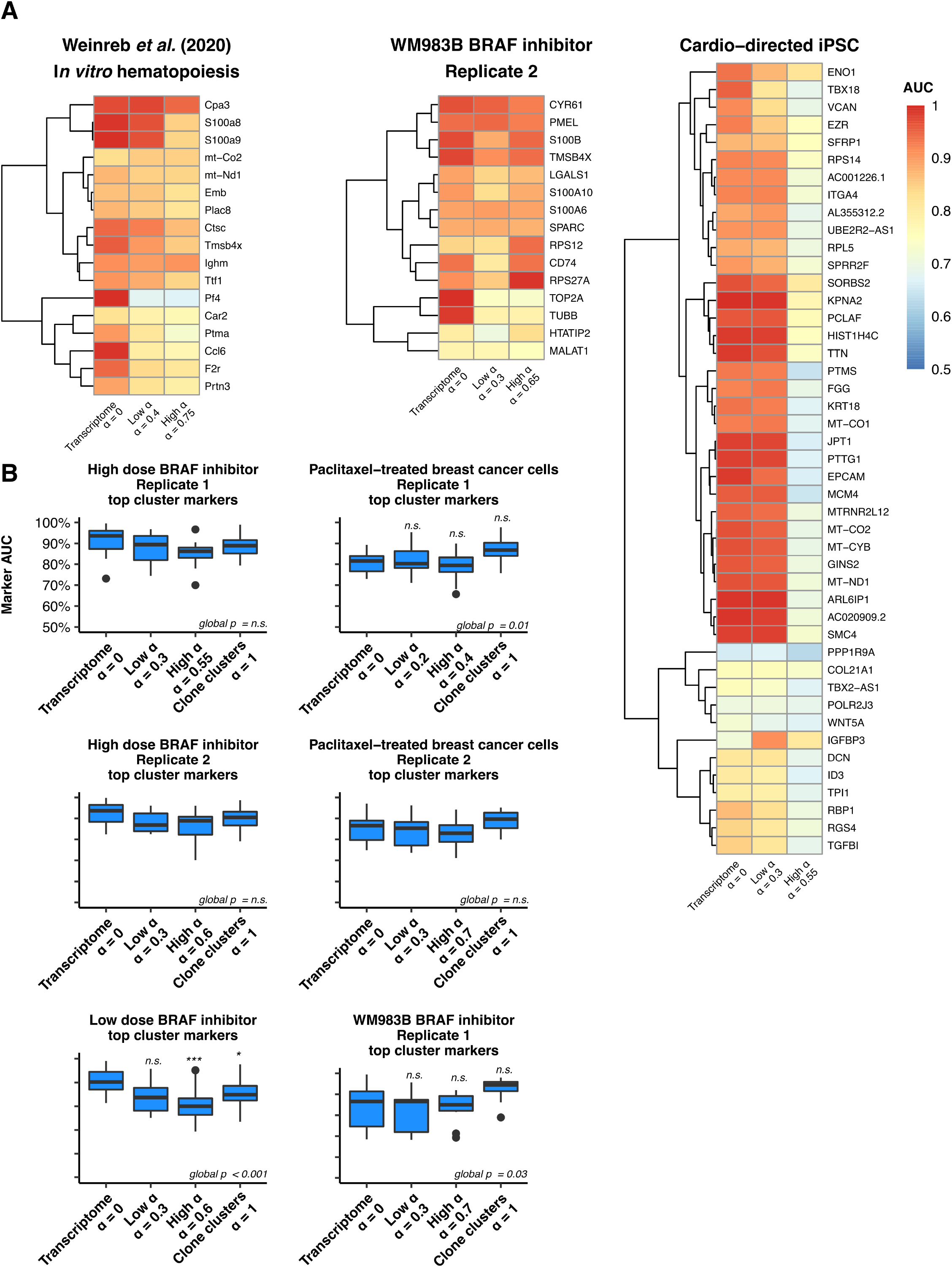
Top cluster marker overall fidelity is preserved across hybrid clusters **A.** Heatmaps of AUC values for the union of all top cluster markers at the transcriptome, low α, and high α level. **B.** Distributions of AUC values for the top marker per cluster at zero, low, high, and maximum (α = 1) values of α for the remaining samples not depicted in Figure 2D (see Methods). “Global p” indicates the p-value for the non-parametric Kruskal-Wallis test with Bonferroni correction. When “global p” was less than <0.05, pairwise Wilcox tests were performed with Bonferroni correction. Annotations above boxes indicate the corrected Wilcox test p-value compared to α = 0 (n.s. - not significant, * - p < 0.05, *** - p < 0.001).

**Figure S6:**
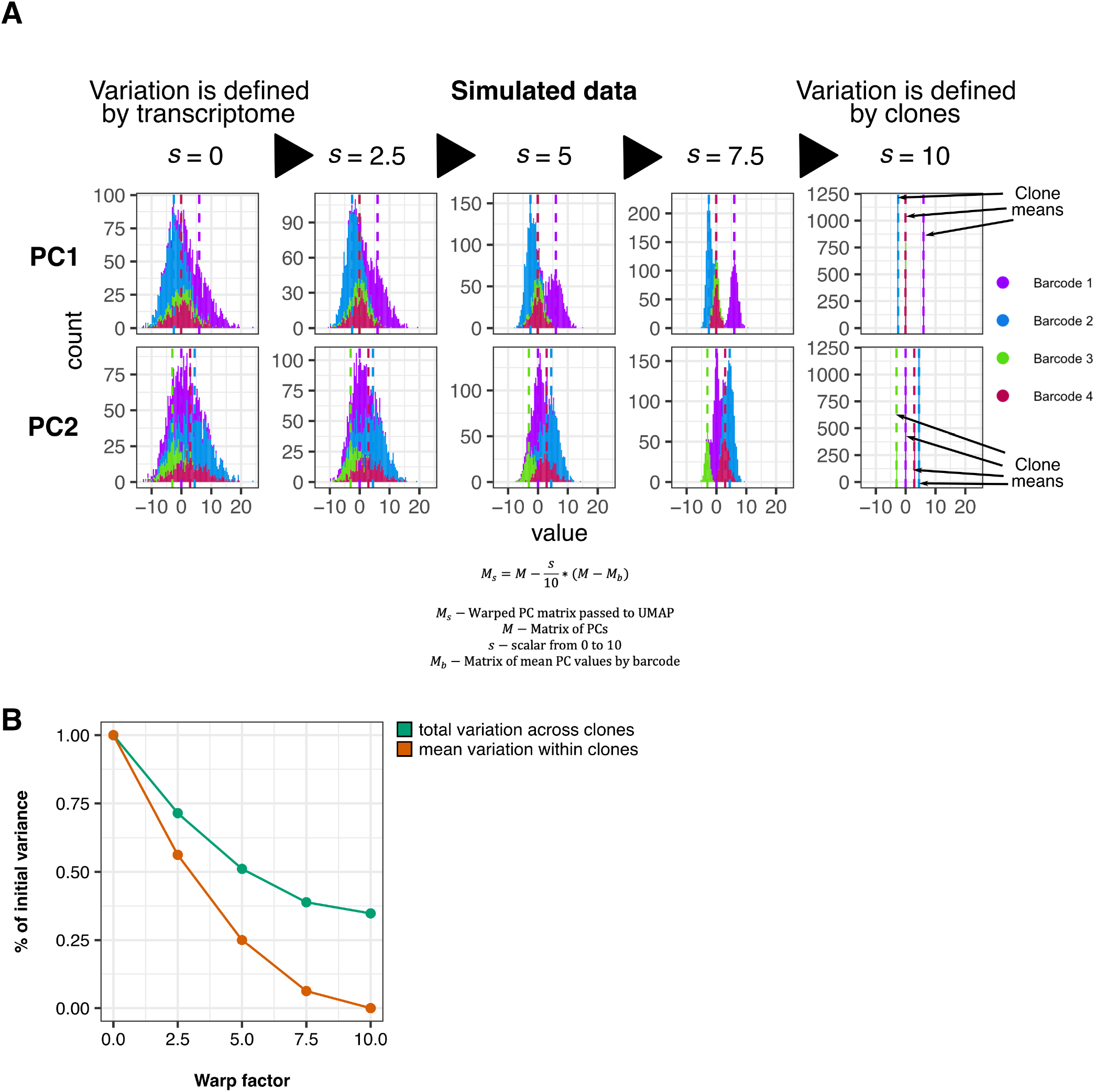
Increasing Warp Factor, *s*, reduces the variation of principal component values around clone barcode means. A. Histograms colored by clone barcodes demonstrating the distribution of PC values for two principal components for simulated data for 3,000 cells with four clone barcode assignments. Vertical dashed lines represent the mean values for each clone barcode at *s* = 0, equivalent to the unmodified PC means. At *s* = 10, modified PC values are equivalent to the assigned clone barcode means. Modified PC values are then passed to the UMAP implementation for visualization as in Figure 4. B. Mean variation of all PCs in simulated data within and between clone barcodes stratified by values of *s*.

## Acknowledgements

YG acknowledges support from the Burroughs Wellcome Fund Career Awards at the Scientific Interface, the Jane Coffin Childs Memorial Fund, and the Schmidt Science Fellowship. AR acknowledges support from NIH Director’s Transformative Research Award R01 GM137425, NIH R01 CA238237, NIH R01 CA232256, NIH P30 CA016520, NIH SPORE P50 CA174523, and NIH U01 CA227550.

## Author Contributions

LPR, YG, and AR conceived and designed the project. LPR, YG, and CLJ compiled and preprocessed data. LPR designed, performed, and analyzed all experiments, supervised by AR. LPR and AR wrote the manuscript with input from all authors.

